# Resolving Complexities in Taxonomic Lineages of the Organellar and Nuclear Genomes of *Galdieria* through Comparative Phylogenomic Analysis

**DOI:** 10.1101/2022.10.04.510841

**Authors:** Manuela Iovinella, Sarah C. L. Lock, Jessica M. Downing, Jennifer Morley, Yen Peng Chew, Luke C. M. Mackinder, James P. J. Chong, Georg A. Feichtinger, Peter D. Ashton, Sally James, Daniel Jeffares, Claudia Ciniglia, Seth J. Davis

**Affiliations:** Department of Biology, University of York, Wentworth Way, York YO10 5DD, UK; Department of Environmental, Biological and Pharmaceutical Sciences and Technologies, University of Caserta “L.Vanvitelli”, Via Vivaldi 43, 81100 Caserta, Italy; Phycosera Ltd., Leeds, LS18 4AW, UK; State Key Laboratory of Crop Stress Biology, School of Life Sciences, Henan University, Kaifeng 475004, China

## Abstract

Exploration of life in extreme environments allows the discovery of intriguing organisms with extraordinary biotechnological potential. An example of extreme environments is represented by hot springs, where harsh conditions (pH < 1; Temperature > 50°C; high concentrations of metals) are prohibitive for most living organisms, except for archaea, bacteria and a few eukaryotes like the unicellular red alga *Galdieria*. Phylogenetic analysis based on a few plastid and nuclear genes highlighted the intricate genetic structure of *Galdieria* and the hypothesis of diverging clades within the *G. sulphuraria* species. To resolve enigmatic relationships between lineages, we used plastid, mitochondrial and nuclear genome-scale data obtained from numerous strains from around the world. The resulting phylogenomic analysis identified: i) the divergence of each of the mitochondrial, plastid, and nuclear genomes into the same six clear lineages; ii) the independent evolution of the lineages; iii) the incongruent interlineages relationships between the three genomes. Differential evolutionary pressure between the strains and the genomes were also highlighted by synonymous and non-synonymous substitutions.

## Introduction

The phylogenetic position of the taxonomic class of Cyanidiophyceae has been discussed for several decades, as these green-colored red algae have been a classification enigma [1]. These species are basal to the Rhodophyta lineage and several species hold promise in various biotechnological studies. Morphological and physiological analysis [2–4] were employed to establish three genera (*Galdieria, Cyanioschyzon* and *Cyanidium*) and eight species (*G. sulphuraria, G. daedala, G. partita, G. phlegrea, G. maxima*, C. *merolae, C. caldarium*, and *C. chilense*.

Genus *Cyanidium* comprises a polyextremophilic species, *Cyanidium caldarium* [5] and a mesophilic one, *Cyanidium chilense* [6]. They are characterized by a very simple morphology: round shape, absence of vacuole and only 1 mitochondrion. The dimensions vary between 2 and 6 µm, [5]. They divide asexually through the production of 4 endospores and display typical pigments are chlorophyll a, C-phycocyanin, allophycocyanin and carotenoids [7]. *Cyanidioschyzon merolae* is the only species belonging to the genus *Cyanidioschyzon* [8]; cellular dimensions are lower than those of *Cyanidium* (1,4 µm x 3-4 µm) indicating an oblong shape; they have 1 mitochondrion and 1 plastid with the same pigments of *Cyanidium*. Important feature of *C. merolae* is the absence of the cell wall and the binary fission as modality of cellular division, differing from the other genus of *Cyanidiophyceae* [8].

Species of the Genus *Galdieria* are facultative heterotrophic microalgae, able to use both ammonium and nitrate as nitrogen source [9]. They are equipped with 1 or more vacuoles, 1 mitochondrion and a multilobed plastid. Four species have been reported to belong to this genus: *G. sulphuraria* [5], *G. partita, G. daedala, G. maxima* [3]. This was based on morphological features, like cellular dimensions, number of endospores, number, and shape of plastids in different moments of the cell cycle [3,5]. In the last years, the employment of molecular tools allowed us in the establishment of the species *Galdieria phlegraea* [1,10] and of the newest genus *Cyanidiocoocus* with the species of *Cyanidiococcus yangmingshanensis* [11].

*Galdieria sulphuraria* is a polyextremophilic red alga thriving in thermoacidic (25°C < T < 30°C; 0 < pH < 5) environments with a remarkable metabolic flexibility, being able of growing auto-mixo- and heterotrophically [9,12]. All *G. sulphuraria* strains share common morphological traits including a single cap-shaped chloroplast. *G. sulphuraria* has drawn interest in biotechnology as it has broad metabolic capacity and can survive feedstock conditions that would be toxic to most eukaryotic microorganisms. Establishing the various strains present in stock centers has been confusing given the various species names defined for the genus based on morphology.

At present it is not clear how many taxonomic species groups of *Galdieria* there are. The cellular ultrastructure of *G. sulphuraria* has been used to classify the strains collected in different worldwide geothermal sites [2–5,13]. Lately, however, it was observed that the elementary shape (small round ball) and the simple ultrastructure that characterize *G. sulphuraria* cells do not match the diversity observed at the molecular, biochemical, and physiological levels [1]. Given their long evolutionary history [14], it is intriguing that only a handful of recognized species have survived in this lineage, and it is more likely that the genetic diversity of these organisms is underestimated. The sampling campaigns in different geothermal areas throughout the world increased the molecular knowledge of *G. sulphuraria*, leading to the general idea that these ancient microalgae have been evolving into more lineages than thought [1,13,15,16].

Phylogenetic analysis based on partial Ribulose-1,5-bisphosphate carboxylase/oxygenase gene (rbcL) highlighted the intricate genetic structure of *G. sulphuraria* and the hypothesis of diverging clades within the species [15–17]. The subdivision of the isolates reflected the geographical position of the populations and thus their isolation caused by not-acid thermal environments and the impossibility of long-distance dispersal [18]. As a result, speciation events have been taking place over thousands or millions of years, and there might be more strains, species, or ecotypes to be discovered [17]. With the increased amount of molecular data, Hsieh et al. 2015 established hypothetical species, identified as Operational Taxonomic Units (OTUs), and confirmed the involvement of the habitat heterogeneity in the increment of genetic diversity within *G. sulphuraria* [15]. The genetic distance between the subgroups was confirmed by analyzing the inter-population pairwise genetic distance measure FST. High values of FST indicated a low level of gene flow between *G. sulphuraria* populations and thus a diverging and isolated evolution [16].

Here we resolved the phylogenetic relationship among a worldwide collection of different *Galdieria* strains using whole genome data, improving the analysis from a gene-level to a pan-genome level. To further understand the evolutionary history of *Galdieria*, plastid, mitochondrial and nuclear genes were analyzed separately and compared to the species ones. Short read genomic data from 43 *Galdieria* strains were used to derive the plastid, mitochondrial, and nuclear coding sequences (CDSs), which were concatenated in single alignments and used to infer the species-level phylogeny. Our data support the split of *Galdieria* genus into two main species, *G. sulphuraria* and *G. phlegrea*. Moreover, organelle and nuclear genomes identified six clear evolutionary lineages within *G. sulphuraria*. Finally, an overall and a lineage-level analysis of the synonymous and non-synonymous substitutions were performed to understand the nature of natural selection forces affected the divergence and the paraphyletic evolution of the species.

## Materials & Methods

### Strains and cultures

*Galdieria* strains were obtained from the Algal Collection of University of Naples (www.acuf.net), the Culture Collection of Autotrophic Organisms (https://ccala.butbn.cas.cz/), the Collection of Microorganisms from Extreme Environments, the Institute of Plant Physiology, Russian Academy of Sciences (http://en.cellreg.org/Collection-IPPAS.php), the Culture Collection of Algae at Göttingen University (https://www.uni-goettingen.de/en/culture+collection+of+algae+%28sag%29/184982.html), the Tung-Hai Algal Lab Culture Collection (http://algae.thu.edu.tw/lab/?page_id=42) (Fig. 1,Tab. S1). All strains were isolated by streaking the colonies across agar plates. This isolation procedure was carried out three times for each line, and a single colony was used to make a master stock. Inoculating the final ones in Allen medium pH 1.5 [19], these were respectively cultivated at 37°C under continuous fluorescent illumination of 45 µmol photons·m−2·s−1.

**Fig 1.**
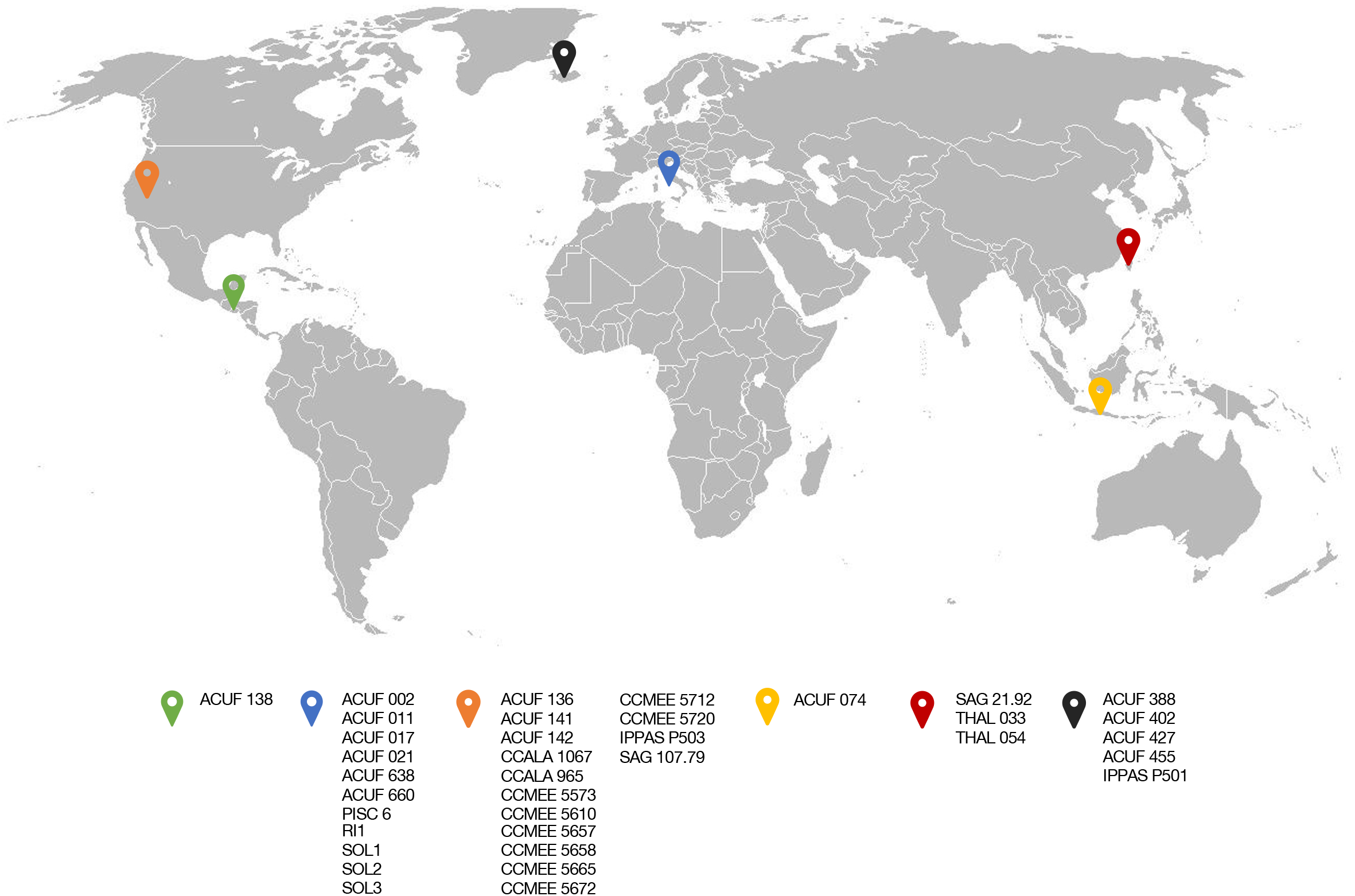
Worldwide distribution of *Galdieria sulphuraria* strains used in this study. Details of the collection sites, along with the sample source and corresponding reference are listed in Tab. S1

### DNA preparation and sequencing

Samples were collected from stock solutions by centrifugation. Total genomic DNAs were respectively extracted using modified SDS-EDTA protocol (Tab. S2) followed by a clean-up step performed with DNeasy Plant Mini Kits (Qiagen, Hilden, Germany). DNA quality and concentration were assessed using a Nanodrop photospectrometer ND-1000 (Thermo Fisher Scientific).

Library preparations were performed with NEBNext® Ultra™ II DNA Library Prep Kit for Illumina Sequencing (Ipswich, Massachusetts, USA), according to the manufacturer’s instructions. Libraries were then sequenced with Illumina MiSeq (Illumina, San Diego, CA, USA) and the resulting reads were trimmed with Trimmomatic [20] and assembled using Spades v3.1 [21]. Assembly quality was assessed using a range of stats shown in Tab S3.

To obtain one single scaffold for plastid and mitochondrial genomes, respective contigs were identified by comparing known proteins from the reference genome, strain ACUF 074 (KJ700460 for mitochondria and NC_024665 for plastid) using TBLASTN from BLAST+ version 2.2.27 [22]. The extracted contigs were re-assembled into one single scaffold using Ragout v. 2.3 and employing the organelle genomes from strain ACUF 074 as the reference [23].

### RNA preparation and sequencing

*G. sulphuraria* cultures (strains 017, 033, 074, 107, 138 and 427) were grown under a 12h/12h light/dark cycle under 42 µmol m^-2^ s^-1^ at 37°C on an orbital shaker (130 rpm). To enhance the diversity of transcripts detected we cultured cells in growth conditions, described in Table S5. Cells were ground into a fine powder with a pestle and a drill in the presence of liquid nitrogen. RNA was isolated and cleaned up using the Monarch Total RNA Miniprep Kit (New England BioLabs, T2010S). RNA quality and concentration were assessed using a Nanodrop photospectrometer ND-1000 (Thermo Fisher Scientific, Waltham, Massachusetts, USA). All RNAs were treated with DNaseI and then pooled by strain relative to the concentration of each sample. RNA quality and concentration were then assessed using the Agilent 2100 Bioanalyzer (Agilent Technologies, Santa Clara, California USA).

RNA library preparation and sequencing were performed at Novogene (UK) Company Limited (Cambridge). Library preparation was performed using NEB Next® Ultra™ RNA Library Prep Kit (NEB, San Diego, CA, USA), employing AMPure XP Beads to purify the products of the reactions during the library prep. Poly-a mRNA was isolated using poly-T oligo-attached magnetic beads, then fragmented through sonication and enriched into 250-300bp fragments. The purified mRNA was converted to cDNA and subjected to the adaptor ligation. The barcoded fragments were finally multiplexed and ran on the Illumina Novaseq 6000 (s4 flow cell) to acquire 20 million read pairs per sample, using the 150bp PE sequencing mode.

### Gene prediction and annotation

Plastid and mitochondrial scaffolds were annotated with GeSeq [24] using *G. sulphuraria*, strain 074W, chloroplast and mitochondrial genomes of the Organelle Genome Resources from the NCBI Reference Sequence Database (RefSeq) as the BLAST reference sequences [25]. Ribosomal RNAs (rRNAs) and transfer RNAs (tRNAs) were annotated using ARAGORN [26] and tRNAscan-SE v2.0 [27]. Finally, the annotated genomes were visualized using OrganellarGenomeDraw v1.2 [28].

Gene prediction was performed on the genome sequences by *de novo* gene prediction using the program AUGUSTUS [29] with parameters trained from *G. sulphuraria* strain 074W, obtained from GenBank (www.ncbi.nlm.nih.gov). Untrimmed RNA sequencing reads were aligned to their respective Illumina assemblies using the STAR aligner v. 2.7.3 [30]. Alignments were filtered with AUGUSTUS v. 3.3.3 filterBAM and converted to AUGUSTUS hints with bam2hints using the defaults [31]. The annotation was performed with AUGUSTUS using both the generated hints and *de novo* gene prediction. Coding sequences with over 94% identity were removed with CD-HIT v. 4.8.1 in order to remove redundant gene sequences [32]. Predicted amino acid sequences were generated with EMBOSS transeq v. 6.6.0 [33] with a minimum ORF length of 40. Sequences without start codons and with stop codons contained in the sequence were removed. All the annotated genomes were deposited into the GenBank database (Tab S6).

### Phylogenetic analysis

Plastid and mitochondrial CDSs were retrieved from the assemblies using tblastn [34] and Geseq [24], using the reference genomes of *G. sulphuraria*, strain 074W. The nuclear gene data set was retrieved from already published sequence data of strain 074W obtained from GenBank (www.ncbi.nlm.nih.gov). Nuclear genes went through a filtering process to use only suitable genes in further phylogenetic analysis. Firstly, all genes associated with the mitochondria and plastid genomes were removed. To identify orthologs of the 074W genes in each of the other genome assemblies, blast databases were created for each of the genomes, then the remaining nuclear genes searched against each of these databases using BlastN from the BLAST 2.10.0 program [34]. The results were assessed using relative blast hit scores where a heat map of scores were clustered by gene (Fig. S1). All genes scoring an average match >0.4 were taken onto the next stage of analysis.

The orthologous CDS sequences for each gene were then aligned using MUSCLE 3.8.31 [35] and the resulting alignments were uploaded to Gblocks v 0.91 b [36] to remove poorly aligned regions applying the options -t = d -b5 = h. For the nuclear alignments, genes that were represented by <40% of the original gene alignment length were excluded; all gene sequences were concatenated by using Gblocks v 0.91 b [36]. The final plastid, mitochondrial and nuclear alignments comprised 13609 bp, 116936 bp, and 5,212,746 bp DNA positions, respectively. Two red algal taxa belonging to *Florideophyceae* and *Bangiophyceae* along with *Cyanidioschyzon merolae*, strain 10D (Cyanidiophyceae) were chosen as outgroup taxa (Tab. S1).

Maximum likelihood (ML) analyses were performed with IQ-Tree v. 1.6.9 [37], using the best substitution model estimated under the partition scheme selected by the program (-spp, -m TEST). Phylogenetic trees were inferred applying 10000 ultrafast bootstrap replicates UFBoot; [38] and 1000 replicates of the approximate likelihood ratio test [aLRT] and Shimodaira-Hasegawa SH-aLRT; [39] for the branch statistical support.

### Consensus Network Analysis and Estimation of Gene Concordance Factors

Analysis of each species tree compared with each other was performed using SplitsTree 5.0.0_alpha [40,41]. The phylogenies of the nuclear, plastid and mitochondrial genomes based on 3532, 124 and 17 genes were produced consisting of 43 taxa. These were imported into SplitsTree to construct a consensus network, where under default options The Consensus Network method was used [42]. Gene concordance factors (gCF) were measured to complement previous phylogenetic analysis. From the concatenation of all 3532, 124 and 17 genes for the nuclear, plastid and mitochondrial genomes each of the corresponding phylogenetic species’ tree were used as the reference trees in the analysis. Each gene tree was also inferred for each locus alignment using IQ-TREE with a model selection. Finally using these trees gCF were calculated using IQ-TREE with the specific option -gcf [43].

### Estimation of non-synonymous to synonymous substitutions ratio

For each of the core species, FASTA format sequences of all protein-coding genes, and their corresponding translations were obtained. In cases where there were two or more transcript variants, the longest transcript was selected to represent the coding region. Protein sequences were aligned using MUSCLE v.3.8.31 [44] and converted into codon aligned nucleotides using PAL2NAL [45]. Nonsynonymous substitutions per nonsynonymous site (dN), synonymous substitutions per synonymous site (dS), and dN/dS (ω) values were calculated for each protein-coding gene using CODEML program in the PAML v 4.3 package [46]. The average dN/dS (ω) were calculated using the M0 model, which calculates the average *ω* for the whole gene, over all branches in the phylogeny. To search for cases of positive selection M7 and M8 models were also used. M7 is defined by using the beta distribution to describe dN/dS variation among sites, where dN/dS value is in the range 0 to 1 (no positive selection is allowed). M8 is the same as M7 except it does allow for positive selection, so some dN/dS sites are >1. M7 and M8 were compared using a likelihood ratio test (LRT) to obtain LRT statistic (twice the difference of the log-likelihood between the null model and alternative model). Here this null hypothesis is that no positive selection is taking place (M7). The resulting list of genes was then filtered for any enzymes with predicted hydrolase function and signal peptides using The UniProt Consortium 2021 and SignalP5.0 [47].

### Comparative genome analysis

Plastid and mitochondrial scaffolds obtained from the re-assembly step were aligned using the ProgressiveMauve algorithm [48] and applying the default settings.

The predicted amino-acid sequences of *G. sulphuraria* proteins were clustered into orthologous groups using OrthoFinder (version 2.3.11) software [49]. Orthologs were identified and clustered by an all-versus-all protein comparison with predicted proteins of the core 6 strains along with extremophile *Cyanidioscshyzon merolae* and mesophilic *Porphyridium purpureum*.

## Results

### General Features of Plastid and Mitochondrial Genomes

Phylogenetic tools were employed to investigate the evolutionary relationships within the extremophile group of *G. sulphuraria* by analyzing and comparing organellar and nuclear protein-coding genes of algal strains. Draft genomes were generated for 43 *Galdieria* lines (Fig 1). Compared with those of the non-*Galdieria* red algae, *G. sulphuraria* organelle genomes are conserved and smaller (13% of reduction for plastid genome and 40% for mitochondrial one). Moreover, plastid genomes showed a reduced GC content, while the mitochondrial ones showed an increase of it. Plastid genomes range between 159 kb and 168 and are constituted by a large single copy (LSC) region, a small single copy (SSC) region and two repeats inverted to each other (IRs, Fig 2). Repeat length vary between strains (Fig. 2). Most of the plastid genomes contain 39 intron-less tRNA genes and 182 protein-coding genes coding for diverse functions connected with the production of pigments, metabolism, photosynthesis, biosynthesis, transcription, and translation machinery (Tab. S4). Unlike the others, strain SAG 107.79 reveals a reduction in both intron-less tRNA (35) and coding genes (167).

**Fig 2.**
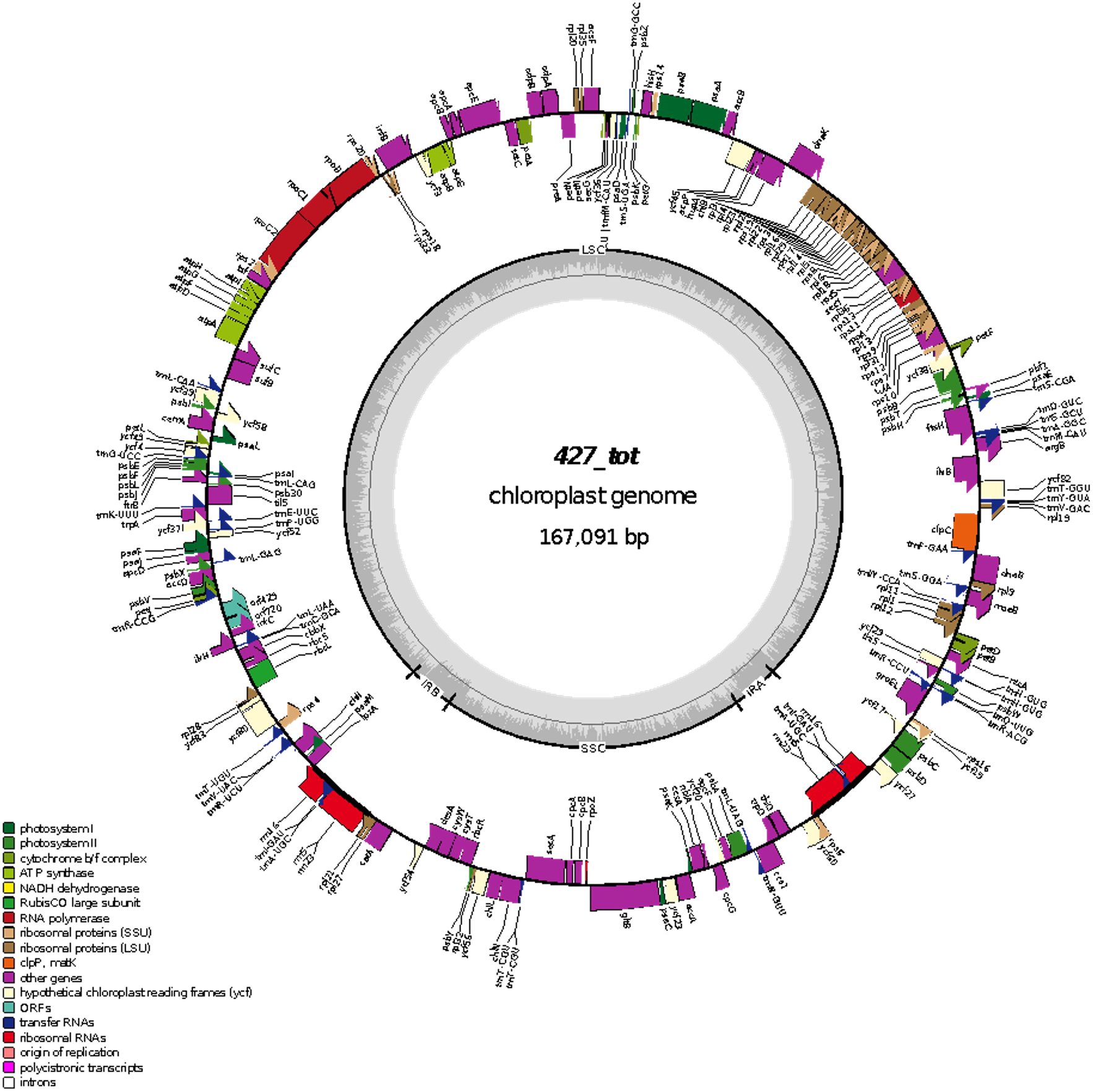
Map of *Galdieria sulphuraria*, strain ACUF 427, plastid genome. Functional categories of the genes are identified with the colored blocks labelled in the legend.

Mitochondrial genomes show a discrete GC content variability ranging from 41.83 to 44.90%. Likewise, genome sizes vary between 20.554 bp and 21.787 bp. An extensive non-coding and variable region is present in all the *G. sulphuraria* mitochondrial genomes. Commonly, all the genomes comprise genes coding for 2 rRNA, 7 tRNA and 19 CDS coding for proteins involved in the respiration system (Tab. S4). In the same area, but on the other strand, some isolates reveal an open reading frame (ORF181), which is highly variable among them (Fig 3).

**Fig 3.**
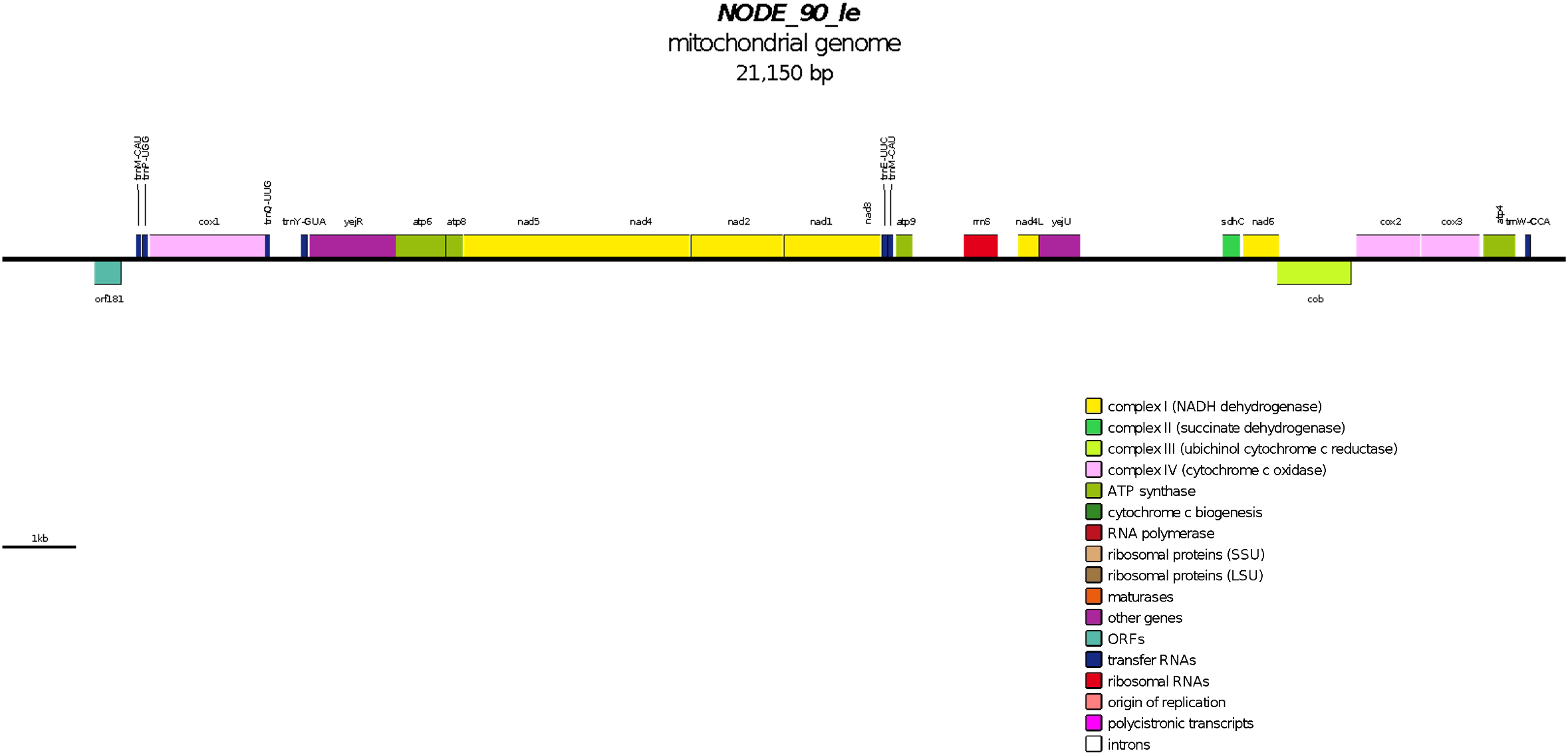
Map of *Galdieria sulphuraria*, strain ACUF 427, mitochondrial genome. Functional categories of the genes are identified with the colored blocks labelled in the legend.

### Phylogenetic analysis

Both organellar and nuclear genome analysis confirmed the monophyletic origin of Cyanidiophyceae (100% UFBoot, 100% SH-alRT). Within the strains identified as *G. sulphuraria*, we identify six lineages that may represent isolated populations each (100% UFBoot; Fig 4). Genome comparisons also highlighted incongruences in the phylogenetic relationship between the six lineages. *G. phlegrea* clearly falls within the *G. sulphuraria* clade. According to plastid phylogenomics, based on 126 concatenated genes, the ancestor organism that originated *G. sulphuraria* assemblage originally mutated, giving rise to the strain ACUF138 (100% UFBoot and SH-alRT). The early divergence of this strain, collected from San Salvador (El Salvador, Central America), caused a high accumulation of mutations, representing 9-10% of the total alignment length compared to all other strains (Tab 1). A following diverging event (100% UFBoot and SH-alRT) generated *Galdieria* strains that colonized the acido-thermal areas surrounding the Mediterranean Sea (Rio Tinto, Italy and Turkey). The strains belonging to this clade derived from a common ancestor and diverged less than 5% (data not shown). Instead, the whole lineage is separated from the others by more than 12000 bp, which is 9-10% of the total (Tab 1). Alongside the evolution of the lineages mentioned above, further diverging events led to the origin of more separate subgroups in *G. sulphuraria*.

**Fig 4.**
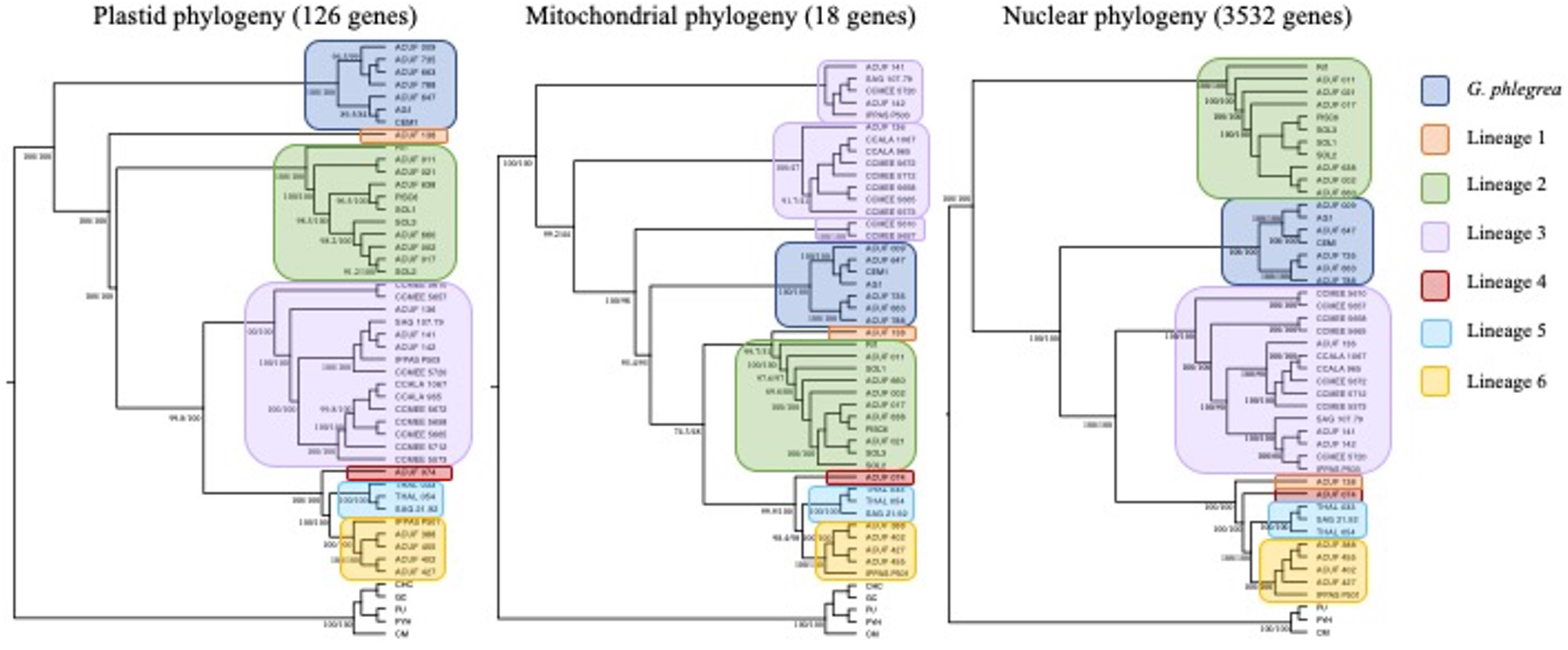
Comparison of the plastid, mitochondrial and nuclear Species trees of Cyanidiophyceae. The phylogenies were inferred from Maximum Likelihood (ML) analysis using the concatenated plastid, mitochondrial and nuclear CDSs, and the partition scheme for the best substitution model. Ultrafast bootstrap (UFBoot) and the Approximate Likelihood Ratio Test [aLRT] and Shimodaira-Hasegawa (SH-aLRT) support values are indicated near nodes. Only bootstrap values ≥ 50% are shown. Different color boxes represent *G. phlegrea* and *G. sulphuraria* lineages and include strains from different geographic locations: blue box = *G. phlegrea*; orange box = San Salvador strain; green box = Mediterranean strains; purple box = strains from the Atlantic region plus US; red box = Java island strain; light blue box = Taiwanese strains; Yellow box = Icelandic and Russian strains; CHC = *C. crispus*; GC = *G. chorda*; PU = *P. umbilicalis*; PYH = *P. haitanensis*; CM = *C. merolae;* detailed collection site information are listed in Table S1

A large clade includes all the strains from the Culture Collection of Microorganisms from Extreme Environments (CCMEE) and the Culture Collection of Autotrophic Organisms (CCALA). Mexican strain ACUF 136 also gathers with all these taxa, but it slightly diverges from them, constituting a sister taxa (100% UFBoot and SH-alRT). Even this sublineage is characterized by low intrapopulation genetic dissimilarity (0-2%) and a high percentage of divergence (8-10%) with the strains of the other sublineages. Remarkably, two strains from CCMEE algal collection, CCMEE 5610 and CCMEE 5657, originally collected from Yellowstone National Park (USA) and Owakudani (JP) respectively, slightly diverged forming a separated sister clade (100% UFBoot and SH-alRT, Tab 1, Fig 4). Concurrently, another ancestral strain, separated from the latter population, led to the paraphyletic development of strains from Iceland, Taiwan, and Java’s Indonesian island (100% UFBoot and SH-alRT). Even if the plastid phylogeny highlighted the separation of these strains in three main groups (100% UFBoot and SH-alRT, Fig 4), the percentage of the mutations among them reaches only 6%, while the separation of the whole clade from the other lineages confirms the percentage of diverging rate of 8-11% (Tab 1).

Mitochondrial phylogenomics, inferred with 18 concatenated genes, highlights three different diverging events resulted in an early separation of all the strains from the American areas (Fig 4). The first segregation occurrence resulted in one monophyletic subgroup (Lin. 3/A) well supported by the statistical analyses (100% UFBoot and SH-alRT, Fig 4). Phylogenetic tree reveals the subsequent separation of strains from CCMEE and CCALA algal collections, along with ACUF 136 (Lin. 3/B). This separation, anyway, is supported only by SH-alRT statistic (44% UFBoot, 99.2% SH-alRT); neither the percentage of dissimilarity among them (1%) confirms the separation of the lineage in two different groups (Tab 2). The third diverging moment separated the remaining two strains from the same lineage (CCMEE 5610 and CCMEE 5657), as occurs in the plastidial genomes (Lin. 3/C; Fig 4; Tab 2). Remaining *G. sulphuraria* strains group in a large clade and cluster as a sister taxa of the other species *G. phlegrea*, which usually happens with plastid markers (95% UFBoot, 95.4 % SH-alRT; Fig 4). Within this group, Mediterranean populations clustered all together in a monophyletic clade, even if small divergences were highlighted at an intrapopulation level. Separation of these strains was supported by 100% of UFBoot and 100% SH-alRT, and the mean percentage of nucleotide divergence is 10%, with higher values (15%) when comparing it with the early diverged American strains (Tab 2). Surprisingly, the mitochondrial phylogeny groups the above population with the strain ACUF 138 from San Salvador. It is to be noted, after all, that this unusual association is supported only by SH-alRT (51% UFBoot, 99.7% SH-alRT) and that the two lineages differed by up to 15% of their whole mitochondrial genomes.

On the other branch of the large clade, the paraphyletic evolution of the strains from Iceland and Eastern Asia, already highlighted by the plastid genome, was confirmed by the mitochondrial one (100% UFBoot, 99.9% SH-alRT). Strain ACUF 074 confirms a slight separation from the others (up to 13%), while Taiwanese and Icelandic strains confirmed a close relationship by both statistics. These strains were characterized by a very low intrapopulation variation (≤ 1 %) and differed by 7% from each other (Tab 2).

Of the 6851 nuclear genes in *G. sulphuraria* 074W retrieved from GenBank, after initial analysis, 3532 genes had suitable coverage across all 43 genomes to be used to determine the species phylogeny. We observed a highly conserved clade containing mostly Italian strains, Lineage 2 (RI1, 011, 021, 017, PISC 6, SOL1, SOL2, SOL3 638, 0022, 4512) (100% UFBoot and SH-alRT). The whole lineage is separated from the others by around 22-29%. The clade grouping populations of the acido-thermal areas around the Mediterranean Sea (Rio Tinto, Italy and Turkey) shared a common ancestor with *G. phlegrea* (100% UFBoot and SH-alRT; Fig 4; Tab 3).

Further diverging events led to the origin of more separated subgroups in *G. sulphuraria*. The biggest clade included all the strains from CCMEE and CCALA algal collection, Lineage 3. This lineage is characterized by low intrapopulation genetic dissimilarity ∼5% (data not shown) and a high percentage of divergence (16-29%) with the strains of the other lineages. The next divergent event originated the strain ACUF138 (San Salvador), Lineage 1 (100% UFBoot and SH-alRT), with 24-31% dissimilarity with the others. *G. sulphuraria* ACUF 074 (Java Island, Indonesia) forms a single lineage (Lineage 4), (100% UFBoot and SH-alRT), which showed a 23-29% dissimilarity in nucleotide sequence with other lineages. Concurrently, the remaining two lineages were separated from the latter population. This led to the well supported (100% UFBoot and SH-alRT) paraphyletic development of strains collected in Taiwan (Lineage 5) and then Iceland and Russia (Lineage 6). These strains were characterized by an 11% sequence difference between the two lineages and a 16-28% from each of the other lineages (Tab 3).

### Consensus Network analysis and Estimation of Gene Concordance Factors

Concordance factors for each node on the resolved species tree for all genomes were measured and compared with discordance factors, which relate to the proportion of genes that support a different resolution of the node (gDF). The number of gene trees that supported each branching event (gCF) is shown in Fig 5 along with consensus network of the mitochondrial, plastid and nuclear splits tree. The network shows clear distinct separation of groups of strains, further supporting the formation of the six lineages. The concatenated nuclear, plastid and mitochondrial genomes all identify the same six lineages, and the majority of individual genes also support this. The differences in topology that appear in the six lineages between the nuclear, plastid and mitochondrial are also highlighted. This analysis shows all the final six identified lineages are highly supported by the gene trees. Consensus network from all plastid, mitochondrial and nuclear genes highlighted the paraphyletic evolution of *G. sulphuraria* lineages 1, 2, 4, 5 and 6 as already revealed by the species phylogeny, except for Lin. 3. Despite the monophyletic origin of this lineage, strains belonging to this group were subjected to further diversification into small groups (Fig 5). Monophyletic clustering of lineages 1, 2, 4, 5 and 6 is highly supported even by the mitochondrial genes (18 genes over 18 ones used in the concatenated alignment phylogeny). Divergence of the American populations in three different clades, which were statistically supported in the plastid phylogeny as a single monophyletic group, is partly highlighted by the gene phylogenies (18/18 for lineage 3/A, 2/18 for lineage 3/B and 15/18 for lineage 3/C; Fig 5). Gene concordance factors on the concatenated nuclear tree highlighted a significant support for the lineages 2, 5 and 6 (more than 3000 nuclear genes over the total). Lineage 3 shows a discrete support from the individual gene trees (2448/3552), while less than 30% of gene trees support the monophyly of lineages 1 and 4 (Fig 5).

**Fig 5.**
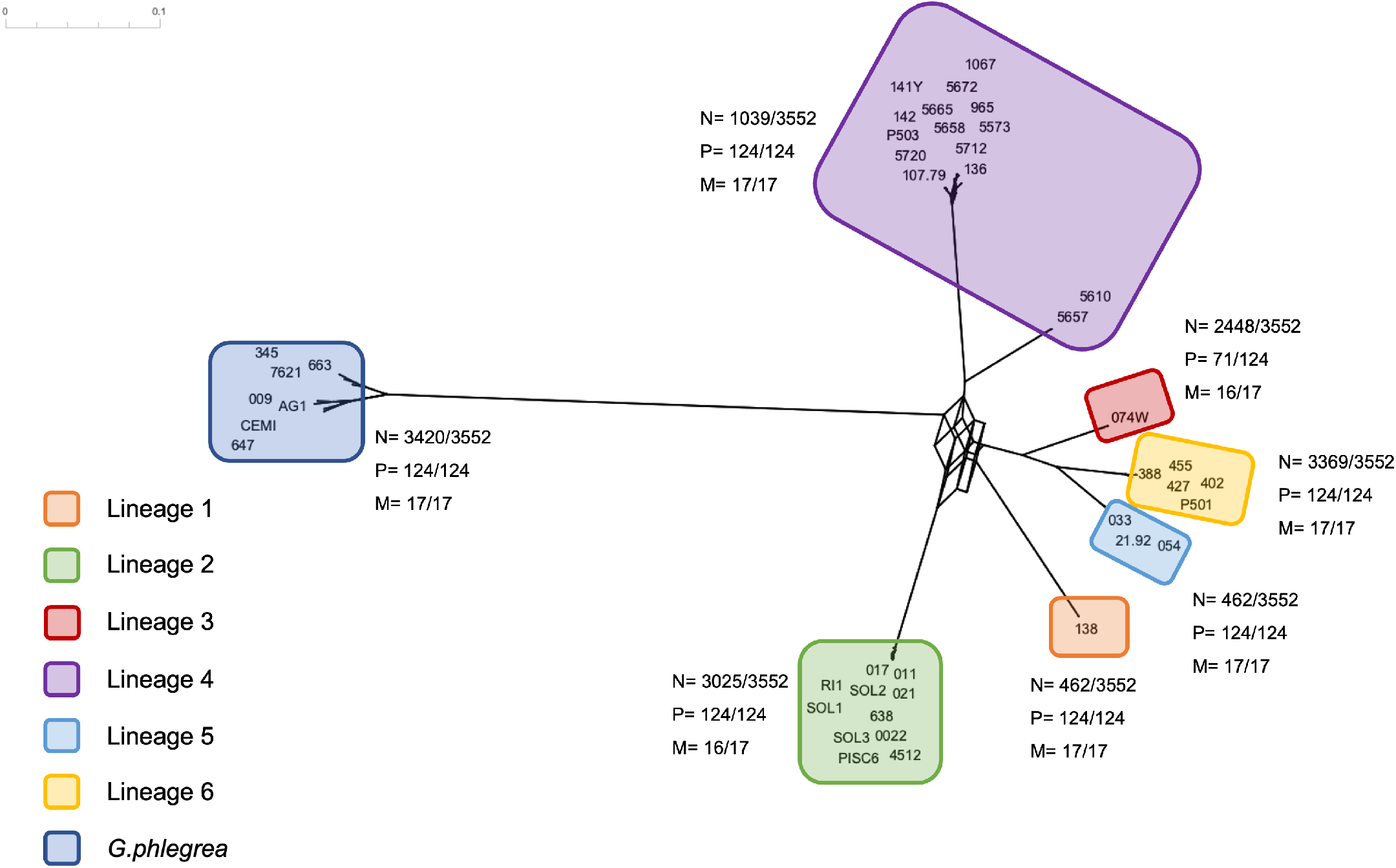
Consensus network of the nuclear, plastid and mitochondrial phylogenies using 43 *Galdieria* strains. The Consensus Network method [38] was used (default options) so as to obtain 116 splits and the Splits Network Algorithm method [39] was used (default options) giving splits network with 290 nodes and 516 edges. The number of nuclear, plastid and mitochondrial gene trees that support each lineage in the species tree topology are indicated near the lineages, collected from gene concordant (gCF) analysis

### Evolutionary rates

As highlighted by the previous analysis, the majority of the divergence observed is between the six lineages and not within them. Therefore one *G. sulphuraria* strain from each of the six lineages identified was selected to represent each lineage in further analysis. These are as follows: 017 (Lin 2), 033 (Lin 5), 074W (Lin 3), 107.79 (Lin 4), 138 (Lin 1) and 427 (Lin 6).

Substitution rates were measured in the plastid, mitochondrial and nuclear genomes between the six *G. sulphuraria* lineages using the core six strains (Table 4). Our analyses included 126 plastid, 18 mitochondrial and 1947 nuclear encoded genes that were present in all six genomes. Nucleotide substitution rates in *Galdieria* are higher in the nuclear genome than in the mitochondrial and plastid genomes. Synonymous site divergence was found to be essentially saturated in mitochondrial, plastid and nuclear genes, with estimates of 1.9, 1.6 and 2.7 substitutions per site respectively (Table 4) meaning that the synonymous divergences is essentially too large to be estimated accurately. Nonsynonymous site divergence is much less with average rates 0.22 (mitochondria), 0.1 (plastid) and 0.36 (nuclear) substitutions/site.

The rates of substitution at nonsynonymous sites (dN) were lower in all three compartments 0.22 (0.22), 0.1 (0.07) and 0.36 (0.25), but were, on average, 1.6 and 3 times greater for the nucDNA relative to the mtDNA and ptDNA. The dN/dS ratio, which can be used to gauge the intensity and directionality of selection, was fairly different across the three genomes. The average for plastid, nuclear and mitochondrial genomes being 0.07 (0.05), 0.13 (0.07) and 0.42 (1.36) which is consistent with the strongest purifying selection acting on the plastid genome. Whilst the mitochondrial genome is presented with the highest average dN/dS ratio when looking at the SD this is also extremely high, this was caused by the *nad6* gene having an above average ratio and skewing the Using the adjusted mitochondrial dN/dS 0.09 (0.08) then substitution rates at non-coding sites were also highest in the nuclear genes. Fig 6 shows the plots of the dN vs dS of the plastid, mitochondrial and nuclear genes. The majority of genes have dN <1 and dS <10. Additionally, Fig 6 shows the distribution of dN/dS = ω to be normal and most genes have ω ≤ 0.2 as is expected. For all genes the ratio of substitutions is never above 0.43. Analysis of the dN and dS values under different models was used to assess any genes under positive selection. This resulted in two mitochondrial genes (*nad2* and *nad6*), three plastid genes (gltB, psaA and ycf80) and 288 nuclear genes (sup x) showing positive selection that would require further investigation.

**Fig 6.**
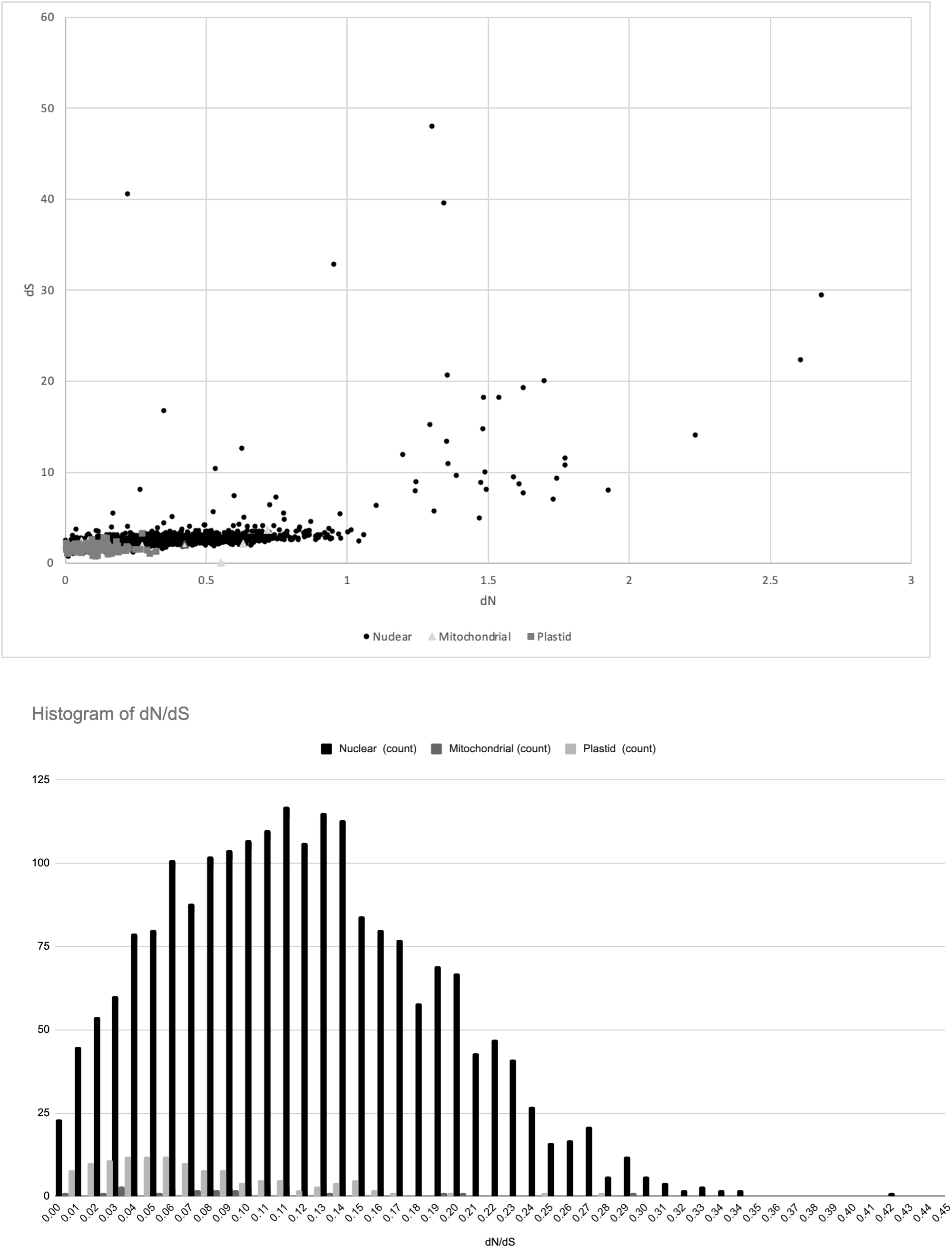
(Top). Pairwise omega (dN/dS) values. This graph shows pairwise dN vs dS values for *Galdieria* nuclear, mitochondrial and plastid genes. The line is dN/dS = 0.5. (Bottom) Histogram showing the distribution of ω for each of the nuclear, mitochondrial and plastid genes.

### Comparative genome analysis

Both plastid and mitochondrial genomes were highly conserved among *G. sulphuraria* species (ACUF 138, ACUF 017, SAG 107.79, ACUF 074, THAL 033 and ACUF 427) representative of the six lineages with some small variations. Analysis of plastid genome synteny identified 23 locally collinear blocks (LCBs) with no or little inter-block spacers, which is consistent with the relatively low sequence divergence described above. LCBs identify conserved segments that appear to be internally free from genome rearrangements. Comparing the genome synteny of *Galdieria* strains with that of *Cyanidioschyzon merolae*, there is a large rearrangement of the gene clusters including translocation and inversion (Fig S2). Inverted gene clusters mainly contain iron-sulfur and sulphate transporters (sufB, sufC), ribosomal proteins (rpl, rps), photosystem II and I proteins (psa, psb). Mitochondrial genomes syntenic analysis identifies mainly 3 LCBs, which are consistent in all the *G. sulphuraria* lineages (Fig S3). The arrangement of the LCBs is also uniform with that of *C. merolae*, except for the block containing the nad6 and cytb genes, which is inverted.

The total number of genes in each of the *G. sulphuraria* genomes varied along with genome size (Table 5). Strain 017 was the largest genomes contained the highest number of genes, while strain 033 contained the lowest number of genes, it was in fact the third largest genome. *C. merolae* has the lowest number of genes but a genome larger than 5/6 of the *G. sulphuraria* strains. The *P. purpureum* genome was of similar size to strain 017 but contained over 3000 more genes. On average across the *G. sulphuraria* genomes GC content was 38% this is lower than the values given from the *C. merolae* (54%) and *P. purpureum* (55%). Cluster analysis on the three species and eight genomes (*G. sulphuraria, C. merolae* and *P. purpureum*), identified 6298 orthologous clusters. Analysis of these clusters suggested that on average across the *G. sulphuraria* genomes 75% of orthogroups contain proteins from the species, compared to 52% from *C. merolae* and 64% *P. purpureum*. Among the set of homologous genes, there were in total 313 single copy orthologs in *G. sulphuraria*. This ranged across the strains with strain 033 containing the lowest species-specific orthogroups (7 genes) to strain 107 with the highest (145 genes). *C. merolae* contained 142 species-specific gene and *P. purpureum* contained 2473 (Table 5).

## Discussion

### General Features of Plastid, Mitochondrial and Nuclear Genomes

Simple morphology of *Galdieria* strains and a small number of unambiguous morphological features makes the identification of different species and genera a challenge when studying these microalgae. This is the case for the classification of *Galdieria*, hence the use of techniques like whole genome seqeucning are indispensable when trying to understand the taxonomy and fundamental biology of the organism.

Initially the sequencing of the 43 genomes discussed in this study revealed a range in size of the genomes, along with initial sequence coverage of genes obtained from reference strain ACUF 074W (ASM34128v1), showing extremely varied coverage across the different genomes. Even when using the not statistically supported nearest point clustering algorithm, clusters of strains are obvious, emphasizing the diversity across the genomes along with the likely diverging evolutionary paths (Kalantari and McDonald, 1983). These clusters have likely derived from dispersal events from an ancestral population. As thermophiles, whose optimal temperatures are 37° C [50], these species likely colonize relatively limited environments, so it is unclear how these dispersal events occurred. As the nonsynonymous divergence between these lineages is considerable, it is likely that these lineages have been separated for a considerable period. For example, Zhang et al 2022 estimated that the two lineages of *Sargassum* brown macroalgae diverged 300,000 years ago [51]. In these lineages the average nonsynonymous divergence (dN) of mitochondrial genes was 0.04 and average dN for plastid genes was 0.006. The *Galdieria* lineage nonsynonymous divergences we describe are far greater; 0.22 for mitochondrial and 0.1 for plastid genes. Unless mutation rates in *Galdieria* are far higher, we can infer that these lineages are very ancient, and dispersal events have not occurred very often.

Eukaryotic genomes vary dramatically in size and gene counts, typically these factors reveal little about the complexity of the organism. However, genome size does matter [52,53]. Genome size is influenced by many things including the rate at which changes in the base DNA occur (deletions and insertions) along with how effectively an organism reacts to these changes and whether they are selected for or against [54]. Genome size is also closely linked to morphology, namely cell size, *i*.*e* typically larger cells will have larger genomes, it is suggested that cell size can explain a high proportion of variation in genome size [55–59]. It has been shown that typically species with larger genomes will have lower metabolic rates, as well as developing and growing at a decreased rate compared to species with smaller genomes [56,59–61]. The correlation between cell size and genome size is based on that an accumulation of redundant DNA (transposons, introns, junk DNA) will have a fitness cost. Meaning that producing excessive or large amounts of DNA is an energetic burden to the cell, it then follows that larger cells will better tolerate larger genomes. A recent study by Malerba et al., 2020 found this to be true for a eukaryotic green alga *Dunaliella tertiolecta*, there was direct evidence that reduction in relative genome size showed associated fitness benefits. In terms of total biovolume and maximum growth rate a higher fitness was observed in lineages that contained relatively smaller genomes [59].

Previous research on sister species *Cyanidioschyzon* that has a similar sized genome revealed that had condensed its genome size by a reduction in the number of genes and had lost nearly all introns [62,63]. Lynch and Conery, 2003 argue that population size is the main driving factor effecting genome size, that an increase in population size is followed by a decrease in cell size thus causing a decrease in genome size [64]. In this scenario low population size leads to an accumulation of slightly deleterious material (such as transposons, more and larger introns), which leads to an increase in cell size. This is as the relative efficacy of purifying selection vs genetic drift is lower when population sizes are lower. This provides an explanation for *Galdieria’s* relatively small genome and suggests the range in genome size seen across the different lineages could be due to separate populations experiencing different circumstances influencing genome reduction, cell size and population size.

A recent study by Xu et al., 2020 showed that plants exposed to high selective pressure of extreme environments caused the independent appearance of the same trait in different lineages (genomic convergence) [65]. Multiple types of conversion events were found, examples included changes in gene copy number, amino acid usage, gene expression, and even GC content [65]. Thus, as would be expected for *Galdieria*, an organism under extreme environments, the genomes, broadly speaking over all eukaryotes are small and sit towards the lower end of the smallest reported eukaryotic genome ∼ 10 Mb [53]. However, across the 43 genomes there was a 17 Mb range in genome size highlighting once again the diversity shown across the species and the effect evolution of isolated populations can have. Often it is non-coding regions that expand or contact like intergenic regions, introns and transposons and could well be what is happening within these genomes.

Organelle genomes were widely used to infer robust phylogenies in order to understand the evolution of algal plastids [66] or find evidence for organelle genome reduction [67] and rearrangement [68]. The employment of organelle genomes relies on the simplicity to handle because of the reduced quantity of data compared to the nuclear ones and at the same time they contain essential genetic information connected to many vital functionalities [68].

Plastid genomes of 43 *G. sulphuraria* strains analyzed in this study were consistent and congruent with previous findings [69]. All of them demonstrated a partition in four distinct zones (LSC, SSC and two inverted repeats) and the presence of the second inverted region reflected a similar structure with the genomes from Glaucophyta, green algae, land plants, and from all eukaryotic algae with red algal-derived plastid genomes [69]. Thus, it is quite possible that this region was present in the common ancestor and was then lost or rearranged in those lineages that diverged afterwards.

Mitochondrial genomes features, such as genome size, number of CDS, synteny and high CG content are shared by all the isolates and consistent with those described in previous works [69,70]. As stated by Jain et al in 2015, the high GC skew could be a consequence of the heterotrophic life of *G. sulphuraria*, which requires an increase of the energy request from mitochondria when it finds itself living endolithically and in the dark [69]. All the mitochondrial genomes diverged only by a long non-coding sequence of which is not known the exact functionality. Non-coding DNA may have important functions in transcriptional and translational regulation or may be the original site of DNA replication [71]. For instance, the longest non-coding and most variable region in animal mitochondrial DNA is identified as Control Region (CR) and comprises a third strand of DNA, creating a D-loop [71]. In vertebrates this region has strand-specific bias and is involved in the asymmetric DNA replication mechanism [70,72].

### Phylogenetic Analysis

The phylogeny work from this study is congruent with published data and confirms the monophyly of *Cyanidiophyceae* species [1,16,73]. Within gene sequences there was a significant variability between nucleotides, thus leading to the well-supported (100% UFBoot and SH-alRT) phylogenetic divergence of *G. sulphuraria* into six lineages. All phylogenies identify the subdivision of *G. sulphuraria* in more diverging lineages. The six lineages are 1) ACUF 138 from San Salvador, 2) Mediterranean clade, 3) American clade, 4) ACUF 074 from Java (Indonesia), 5) Asian clade and 6) Icelandic clade.

The branches in the nuclear phylogenies leading to the each of the different lineages are always very long in comparison to the terminal branches leading to the single strains which are always very short. This indicates a high divergence between each of the lineages but suggests a low genetic diversity within them. All the lineages diverged from each other upwards of 11% when assessing nucleotide sequence dissimilarity (average 23.6 %); these percentages easily fit into the 8-11% range identified as the threshold level for genus assignment in Rhodophyta [11,74]. Though it should be noted that this threshold level was typically used for multicellular algae and in conjunction with morphological criteria and is not to suggest each lineage is a different genus but rather highlight the extensive diversity and divergence. This work gives strong supporting evidence for the six identified lineages to be separate species; however, this cannot be said with certainty without further research into characterizing the lineages based on other traits and assessing whether lineages populations can interbreed.

### Estimation of Gene Concordance Factors

Analysis of the phylogenetic relationships between the six lineages within the mitochondrial, plastid and nuclear species trees in *G. sulphuraria* reveals an incongruent evolution between the three genomes. Though this is not unexpected as for photosynthetic eukaryotes the relative mutation rates among mitochondrial, plastid, and nuclear genomes have been shown to be different between plants, green and red algae [75–78]. Thus, resulting in different evolution between the respective genomes. The difference in evolution across the three genomes in each case still resulted in the same six lineages being identified.

The majority of genes presented a distinct phylogeny, this is as each individual gene is likely to have a unique evolutionary path as a consequence of meiotic division or some other type of recombination. It is known that different genes will evolve at different rates and be under different selection pressures. Recombination will also result in differing tree topologies and make one consistent phylogeny across the genome extremely unlikely; this is standard when looking at eukaryotic genetic data. Ultimately the divergence into the six lineages is generally well supported but the relationship between the lineages in the branching events leading to the lineages is variable. This is evidence in support of these lineages being isolated populations evolving differently, and that within these lineages there are interbreeding populations.

Analysis of single nucleotide polymorphisms (SNPs) within the *G. sulphuraria* lineages showed linkage disequilibrium, which is also indicative of recombination (see companion paper; Downing et al. 2022). Taking all of this into consideration along with the overall well supported species tree (100% UFBoot and SH-alRT), there is high confidence that the *G. sulphuraria* species tree is resolved. Incongruence between Gene and Species trees is also affected from Gene duplication, hybridization and recombination, all of which are important events contributing to the genetic variation in populations [79,80]. Here there is the production of a second copy of gene, which evolves independently from the original one [80]. The new copy could become a pseudogene or could evolve a new and diverged function, thus creating the phylogenetic coexistence of more gene lineages [79]. Further biological mechanisms inducing phylogenetic incongruence are the incomplete lineage sorting and the horizontal gene transfer [79,80]. The first one implies the persistence of ancestral polymorphisms during subsequent speciation events in a short period of time [81]. Rapid species radiation and the sharing of the ancestral polymorphisms across the strains could result in phylogenetic discordance [81,82].

### Estimation of non-synonymous to synonymous substitutions ratio

High variability of the dN/dS ratios across all genes would suggest that the organelle and nuclear genes could be subjected to different evolutionary forces (Fig 6). It is known by literature that the plastid genes are not tightly linked and undergo to the evolutionary pressure independently. Moreover, dN/dS values, calculated for each pair of lineages and expressed as the mean values of all the genes (Fig 6; Tab. 4), demonstrate a stronger purifying selection for plastid genomes (0.07) than the mitochondrial and nuclear ones (0.42 and 0.13). The evolution of a protein coding gene is influenced by many factors, with correlations to intron number, gene expression and the essentiality of the gene to name a few [83–86]. The types of substitutions acting on the sequence are important, synonymous substitutions within a protein are random and will likely be tolerated across generations. Non-synonymous substitutions are due to neutral evolution and are more often removed by purifying selection, however, a proportion is fixed as a result of positive selection and thus increasing the rate of protein evolution.

Analyzing the rate of the substitutions occurring in a protein can identify information about which selective pressures are happening [87]. The calculation of dN/dS can therefore help to identify genes that are under particular biochemical or ecological constraints, or conversely putative proteins involved in survival adaptation. Analysis of the synonymous and non-synonymous substitutions of the nuclear genes present in all core six strains confirmed different evolutionary pressures across genes. High substitution rates along with events such as horizontal gene transfer (HGT), could be the main evolutionary forces shaping the divergence of the *G. sulphuraria* species, these biological mechanisms have been linked to inducing phylogenetic incongruence [80,88,89]. HGT events have been widely confirmed in *G. sulphuraria* [69,90–92], this may strongly influence the phylogenetic relationship among strains. Genes acquired horizontally are likely to be involved in the adaption of *Galdieria* to its harsh environment, this will include osmotic resistance, salt tolerance, carbon and amino acid metabolism, metal and xenobiotic resistance/detoxification non-metabolic and uncertain functions [92].

High variability of the synonymous substitutions across nuclear genes is generally not considered deleterious as these mutations are typically regarded as neutral or at least have a much smaller effect on fitness, compared to non-synonymous substitutions [93,94]. Calculation of the coefficient of variation (SD/mean; Table x) gives a standardized measure of the dispersion of the distribution of dN to dS, and in this case for each of the genomes dN is much higher than originally calculated and so relatively more variable [95–97]. This could be due to different selection pressures on genes, for example typically proteins such as histones are among the most conserved proteins [98,99] whereas membrane and exported proteins tend not to be [100–102]. It is not unusual in these types of analysis to see dN/dS rates < 0.5 as it is expected that most genes will be under purifying or neutral selection [95]. This is as nonsynonymous changes are more likely to have a functional consequence and therefore will generally be deleterious. This means they are removed from populations more rapidly and thus their rate is typically slower than the rate of synonymous changes. However, a dN/dS <1 does not mean that all genes are under purifying or neutral selection. Genes under adaptive evolution are favoured for and in the case of *G. sulphuraria* can contribute to its adaptation and survival in extreme environments. The analysis for identifying nuclear genes under positive selection revealed 288 genes. These genes are a valuable resource in investigating the adaptations of *G. sulphuraria* and how it not only survives but thrives in low pH and high temperatures.

## Conclusions

This study aimed to understand the phylogenetic relationship among the genus *Galdieria* and different *G. sulphuraria* species. This provided insight into the evolutionary history of the species between their mitochondrial, plastid and nuclear genomes using draft genome sequencing data. It was found that even if morphological traits slightly vary between *Galdieria*, molecular tools allowed the identification of huge variability between them. The resulting phylogenetic analysis identified the divergence of each of the mitochondrial, plastid and nuclear genomes into the same six clear lineages and that these have been evolving separately. Analysis of the synonymous and non-synonymous substitutions confirmed the differential evolutionary pressure between the strains and the genomes and gave rise to multiple genes under positive selection. These genes present good candidates for exploration into *G. sulphuraria’s* adaptation to its environment.

## Supporting information

Support Tables_Tables

## Supplementary Figures

**Fig S1.**
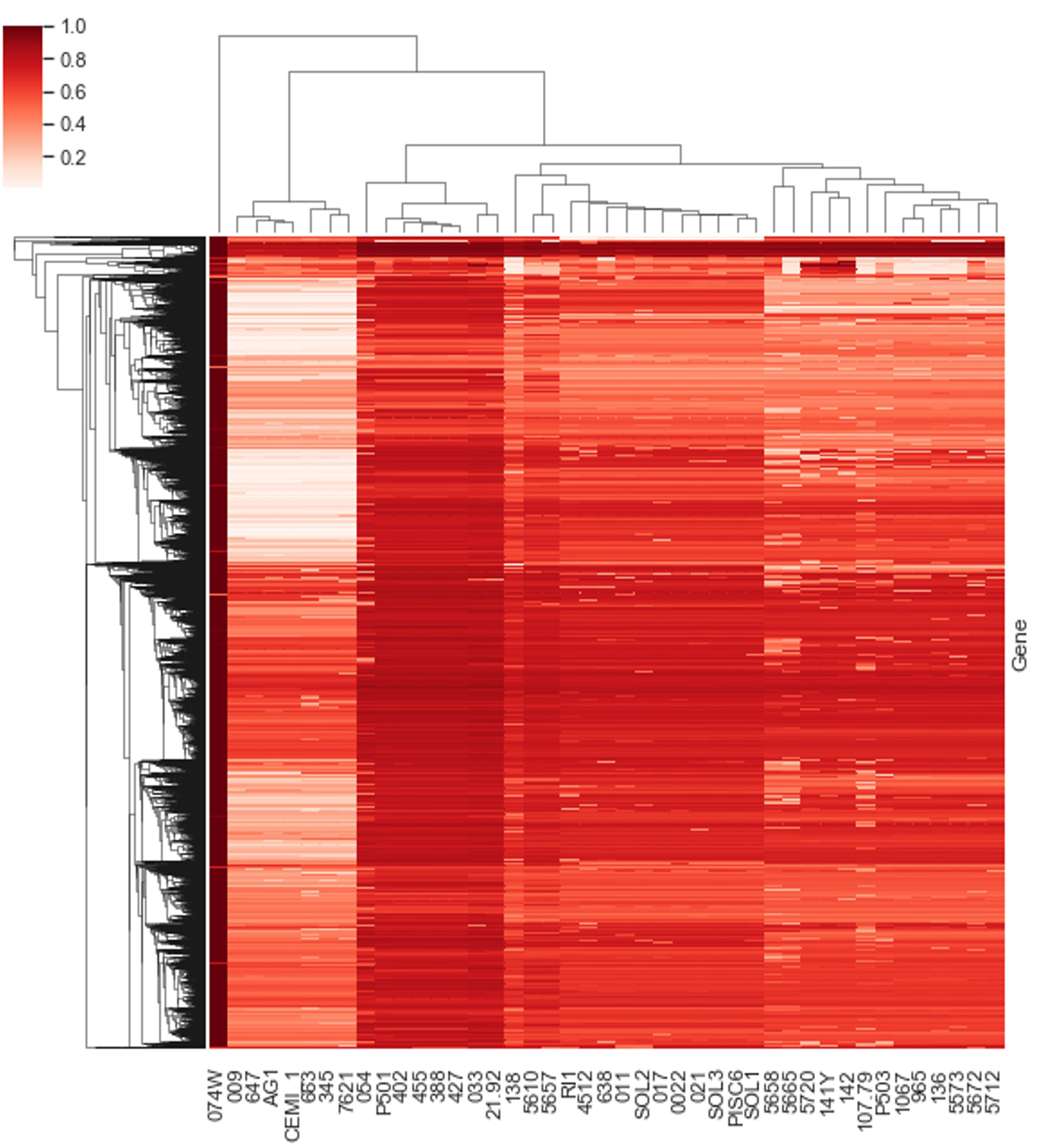
Clustered heat map of all genes scoring an average of 0.4 or greater relative match to reference sequence (percentage identity). Clustered by both Gene (y-axis) and Strain (x-axis) using the nearest point algorithm (Kalantari and McDonald, 1983).

**Fig S2.**
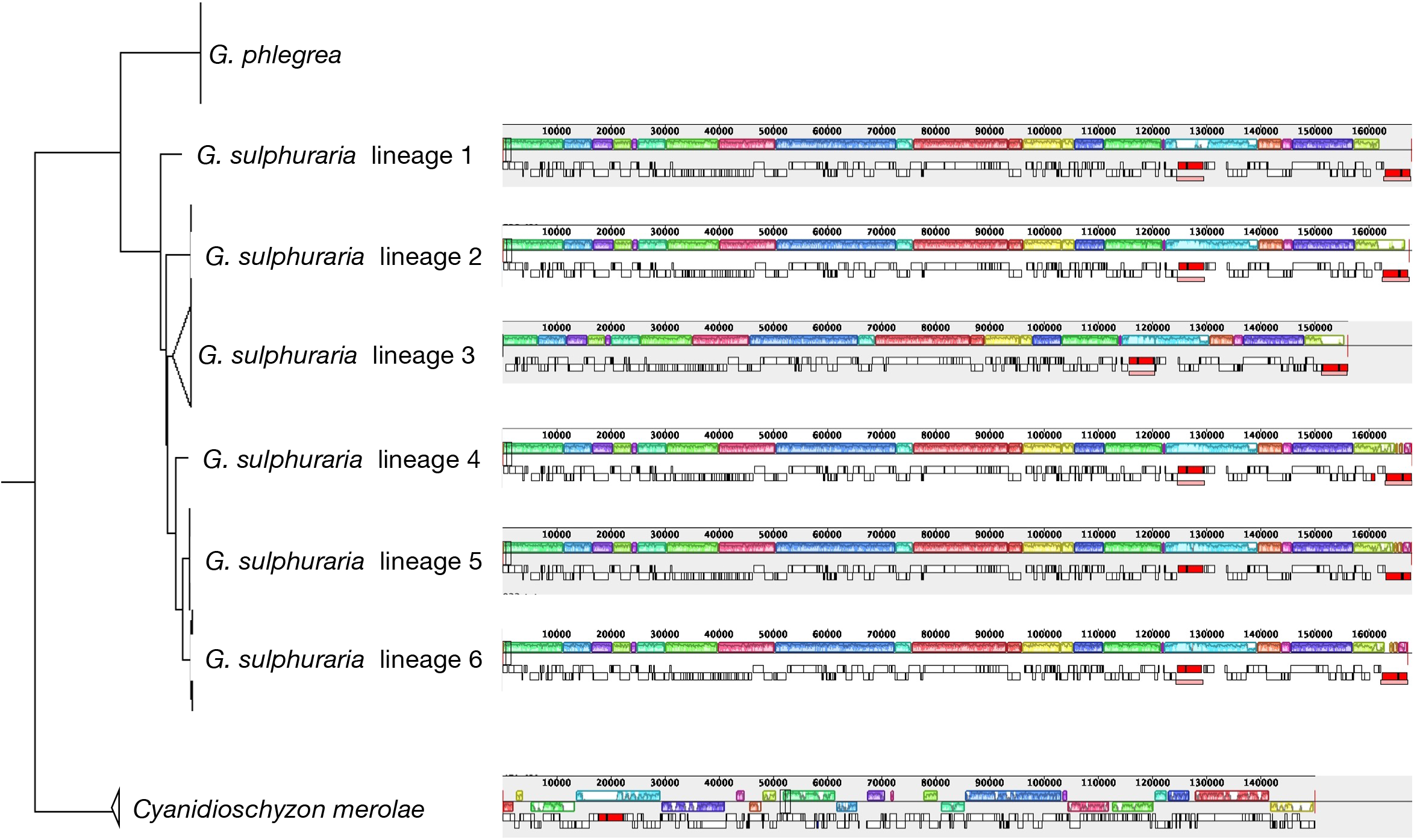
Synteny comparison of *Galdieria sulphuraria* plastid genomes. One representative of each lineage was analysed and compared with the reference genome of *Cyanidioschyzon merolae*. Homologous gene clusters are identified as Locally Collinear Blocks (LCBs) and visualized by the colored boxes.

**Fig S3.**
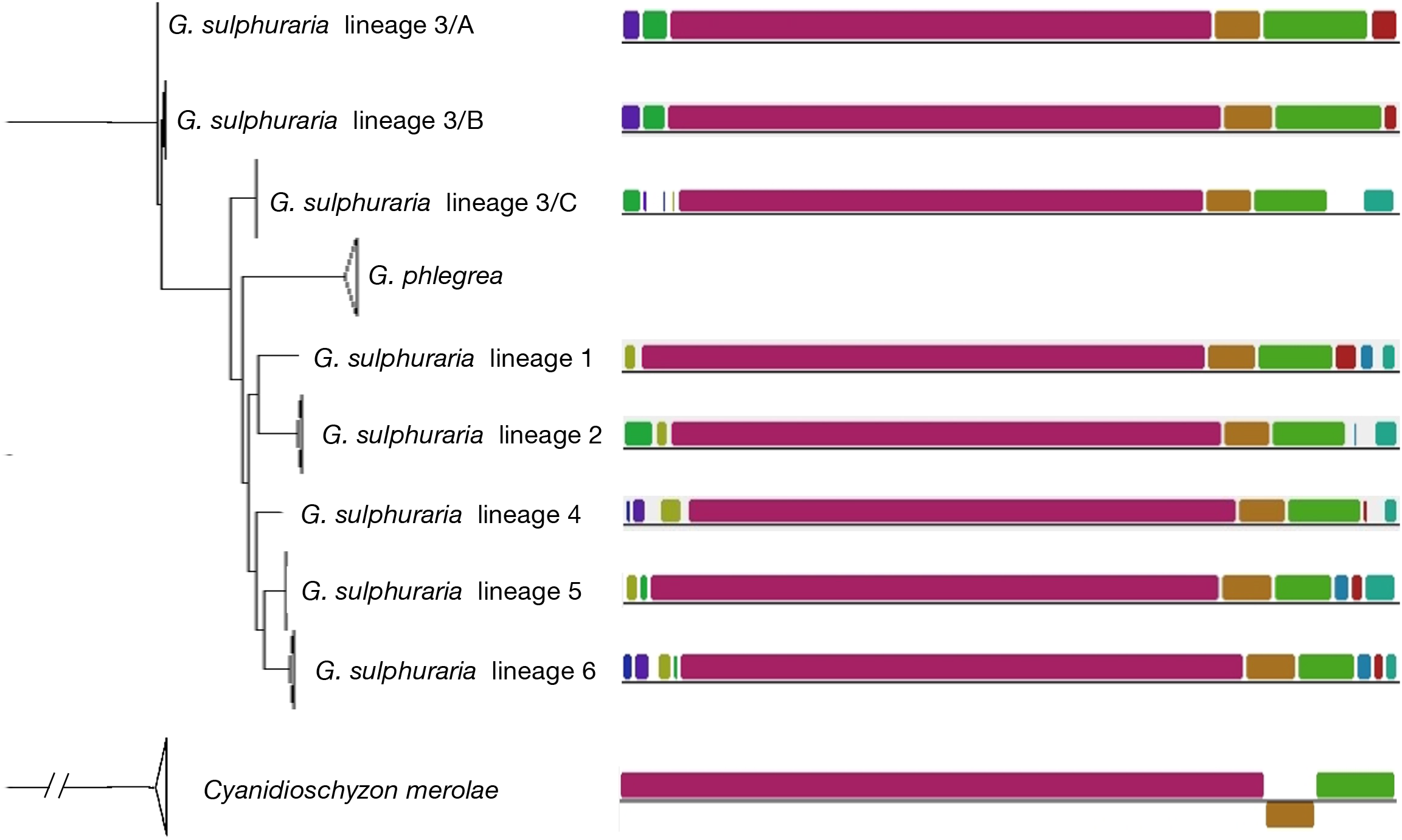
Synteny comparison of *Galdieria sulphuraria* mitochondrial genomes. One representative of each lineage was analysed and compared with the reference genome of *Cyanidioschyzon merolae*. Homologous gene clusters are identified as Locally Collinear Blocks (LCBs) and visualized by the colored boxes.

